# Immunogenicity, safety and efficacy of sequential immunizations with an SIV-based IDLV expressing CH505 Envs

**DOI:** 10.1101/2020.03.06.980680

**Authors:** Blasi Maria, Negri Donatella, Saunders O Kevin, Baker J Erich, Stadtler Hannah, LaBranche Celia, Mildenberg Benjamin, Morton Georgeanna, Ciarla Andrew, Shen Xiaoying, Wang Yunfei, Rountree Wes, Balakumaran Bala, Santra Sampa, Haynes F Barton, Moody M Anthony, Cara Andrea, Klotman E Mary

**Affiliations:** Department of Medicine, Duke University School of Medicine, Durham, NC, USA; Duke Human Vaccine Institute, Duke University School of Medicine, Durham, NC, USA; Department of Infectious Diseases, Istituto Superiore di Sanità, Rome, Italy; Department of Surgery, Duke University School of Medicine, Durham, NC, USA; Beth Israel Deaconess Medical Center, Boston, MA, USA; Department of Pediatrics, Duke University School of Medicine, Durham, NC, USA; National Center for Global Health, Istituto Superiore di Sanità, Rome, Italy

## Abstract

A preventative HIV-1 vaccine is an essential intervention needed to halt the HIV-1 pandemic. Neutralizing antibodies protect against HIV-1 infection in animal models, and thus an approach toward a protective HIV-1 vaccine is to induce broadly cross-reactive neutralizing antibodies (bnAbs). One strategy to achieve this goal is to define envelope (Env) evolution that drives bnAb development in infection and to recreate those events by vaccination. In this study we report the immunogenicity, safety and efficacy in rhesus macaques of an SIV-based integrase defective lentiviral vector (IDLV) expressing sequential gp140 Env immunogens derived from the CH505 HIV-1-infected individual who made the CH103 and CH235 bnAb lineages. Immunization with IDLV expressing sequential CH505 Env induced higher magnitude and more durable binding and neutralizing antibody responses compared to protein or DNA ^+/-^ protein immunizations using the same sequential envelopes. Compared to monkeys immunized with vector expressing Envs alone, those immunized with the combination of IDLV expressing Env and CH505 Env protein demonstrated improved durability of antibody responses at six month after the last immunization as well as lower peak viremia and better virus control following autologous SHIV-CH505 challenge. There was no evidence of vector mobilization or recombination in the immunized and challenged monkeys. Our results show that while IDLV proved safe and successful at inducing higher titer and more durable immune responses compared to other vaccine platforms, the use of non-stabilized sequential envelope trimers did not induce broadly neutralizing antibody responses.

## Introduction

Vaccine induced immunity is generally provided by the elicitation of protective and durable antibody responses. The HIV-1 envelope (Env) is the only target of neutralizing antibodies (nAb)^1^. Several strategies have been evaluated to induce broadly neutralizing antibodies (bnAbs), including the use of various immunization strategies and Env-based immunogens. However, none of these strategies have been effective at inducing broadly neutralizing antibodies (bnAbs) in humans or in non-human primates (NHPs)^2-4^. Antibody-virus co-evolution studies performed on HIV-1 infected individuals from the time of transmission to bnAb development have demonstrated that bnAbs arise after extensive Env diversification^5-7^. Thus, one strategy to induce bnAbs is to perform sequential immunizations with a series of Env isolates from an individual that made bnAbs to mimic natural infection by vaccination and guide bnAb development.

The other crucial aspect of a successful vaccine strategy is the selection of an antigen delivery system that can provide high magnitude and durable immune responses. In the modestly effective RV144 trial^8^, the estimated vaccine efficacy dropped significantly between 6 months to 12 months after vaccination, which correlated with the rapid decline of vaccine induced antibody responses highlighting the importance of maintaining adequate concentrations of protective antibodies over time.

We have shown, in both mice and macaque studies, that integrase defective lentiviral vectors (IDLV) provides prolonged antigen expression and can induce potent and durable antigen-specific immune responses^9-13^. We demonstrated that immunization with an SIV-based IDLV expressing an HIV-1 envelope (Env) induces continued antibody affinity maturation up to three months post-prime, which is further improved upon additional IDLV-Env immunizations^12^. Furthermore we demonstrated persistent antigen expression at the site of injection up to three months post intramuscular immunization^12^, (*Lin et al., 2020* in press) suggesting a direct correlation between the continuous antigen expression by IDLV and the durability of antigen specific immune responses (*Lin et al., 2020* in press).

The SIV-based IDLV has been engineered with a number of features to enhance its safety, including the deletion of the promoter/enhancer elements present in the U3 region of the LTR^14^. This self-inactivating (SIN) feature, together with the episomal configuration of IDLV, markedly reduces the risk of vector recombination that could result in production of replication competent lentivirus (RCL), insertional mutagenesis, and mobilization of the vector from transduced target cells by subsequent infection with a replication competent virus. However, the possibility of recombination between IDLV and the replication competent virus has never been tested in non-human primates.

Here we report that repeated sequential immunization with IDLV expressing gp140 Env immunogens derived from the CH505 HIV-1 infected individual who made the CH103 and CH235 CD4 binding site bnAb lineages^5, 7, 15^, was immunogenic and safe in rhesus macaques. We demonstrate that IDLV-CH505-Env ^+/-^ CH505 Env protein induced higher magnitude and durability of antibody responses compared to protein alone, DNA and DNA+ protein immunization strategies using the same envelope immunogens. Following repeated low-dose intrarectal challenges with the autologous SHIV.CH505.375H.dCT^16, 17^ virus, one out of 8 animals in the IDLV + protein group resisted infection and this group of animals had 3.6 times lower peak viral load compared to the control group of animals immunized with an IDLV expressing GFP (n = 11). Differently from animals immunized with IDLV-CH505 Env alone and animals in the control group, we observed rapid and sustained viremia control in the IDLV + protein group that correlated with the magnitude and quality of the antibody responses. Finally, to evaluate the safety of sequential IDLV-CH505 Env immunization a group of 4 vaccinated animals was challenged intravenously with SHIV.CH505.375H.dCT to enhance the possibility of potential recombination and/or mobilization events between the challenge virus and the IDLV vaccine. No recombination/mobilization events were detected in these animals. Our study demonstrates that IDLV is a safe and effective vaccine platform that induces high magnitude and durable immune responses against HIV-1.

## Results

### Study design

The set of CH505 Envs used in this study were selected based on binding to stages of the CH103 bnAb lineage: the CH505 transmitted founder (T/F) and three natural CH505 variants (week 53, 78, and 100)^15^. Thirty-one Indian rhesus macaques were divided into four immunization groups as shown in **figure 1a**; one group of eight animals was immunized intramuscularly with IDLV expressing sequentially evolved CH505 gp140 envelopes (T/F, w53, w78, w100) at six months intervals (Group B1 – IDLV-CH505 alone). The second group of eight animals was immunized intramuscularly with IDLV expressing sequentially evolved CH505 envelopes in combination with the corresponding CH505-gp140 protein in GLA-SE^18^ adjuvant (Group B2 – IDLV-CH505 + protein). For the last two immunizations of both B1 and B2 group animals we used IDLV expressing a stabilized gp140 SOSIP Env trimer (CH505.w136 SOSIP) with the goal of focusing antibody responses towards the epitopes targeted by different bnAbs^4, 19, 20^. A third group of eleven animals served as the control group and received sequential immunizations with IDLV-GFP (Group B3). At the end of the scheduled vaccinations, monkeys in groups B1, B2 and B3 received ten intrarectal challenges with SHIV.CH505.375H.dCT, whereas, a fourth group of four animals, also immunized intramuscularly with IDLV expressing sequentially evolved CH505 envelopes (Group A), was challenged intravenously to maximize the chances of possible recombination between IDLV and the replication competent SHIV-CH505, as assessed in the safety studies described below.

**Figure 1.**
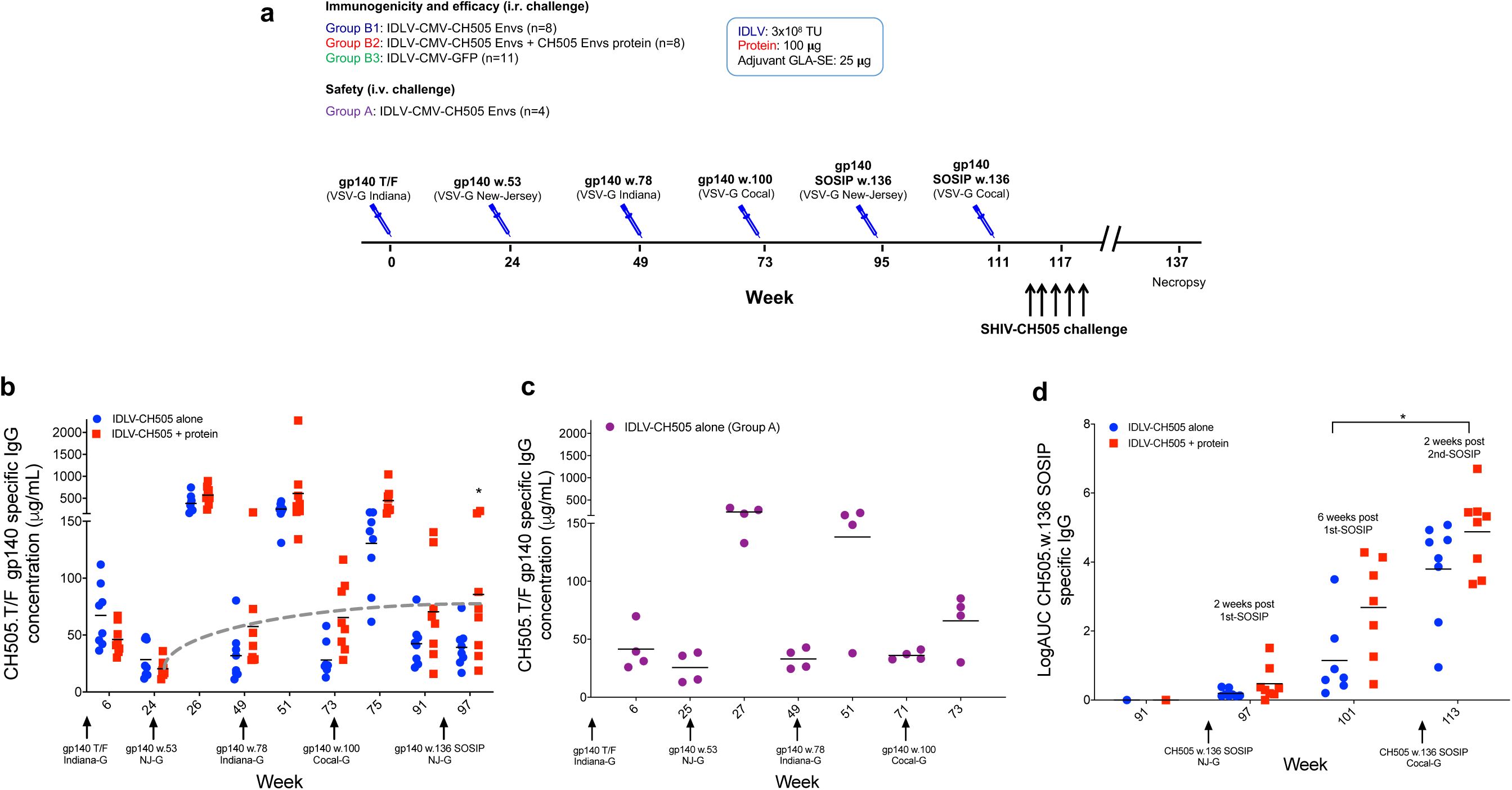
Magnitude and durability of antibody responses induced by sequential IDLV-CH505 Env immunizations. **(a)** Non-human primate immunization regimen with sequential IDLV-CH505 Env ^+/-^ protein. The dose of IDLV, protein and adjuvant used for each immunization as well as the challenge and necropsy schedules are indicated. A group of eleven animals were sequentially immunized with IDLV-GFP as the control arm. (**b**) ELISA binding of plasma antibodies to CH505 T/F Env at the peak and six months post-each immunization with IDLV-CH505 (group B1 animals) and IDLV-CH505+protein (group B2 animals). Binding titers measured as concentration in **μ**g/mL starting at a 1:3000 plasma dilution. The dotted gray line indicates the improved durability of antibody responses at six months post-each IDLV-CH505 immunization. Asterisks indicates that a statistically significant improvement in the magnitude of Ab responses was detected between weeks 24 and 97. (**c**) ELISA binding of plasma antibodies to CH505 T/F Env at the peak and six months post-each immunization with IDLV-CH505 (group A animals, safety arm). (**d**) ELISA binding of plasma antibodies to CH505w.136 SOSIP Env post- IDLV-CH505w.136 SOSIP immunization. Binding titers measured as Log area under curve (Log AUC) starting at a 1:30 plasma dilution.

### Improved durability of Env-specific Ab responses post sequential IDLV-Env immunization

Plasma antibodies (Abs) specific for CH505.T/F-Env were assessed at two weeks post each immunization and then monthly thereafter. All NHPs developed high titers of CH505.T/F-gp140 Env-specific Abs at 6 weeks post-prime (**figure 1b**), that were strongly boosted after each subsequent IDLV-CH505 Env immunizations. Immunizations with IDLV-CH505 alone elicited similar antibody responses as IDLV-CH505 + protein at the prime, however after the subsequent boosts antibody titers were higher in the IDLV-CH505 + protein co-immunization group (**figure 1b**). In this group of animals, antibody titers at 6 months after the IDLV-CH505w.100 gp140 immunization (week 97) were significantly higher (p = 0.0234 based on a Wilcoxon signed-rank test) than those detected at 6 months after the first immunization (week 24) (**figure 1b**), suggesting that the durability and magnitude of the vaccine-induced antibody response improved over time. Similar antibody titers as in group B1 animals were induced in the four animals in group A, also immunized intramuscularly with IDLV-CH505 alone (**figure 1c**).

We observed that following the fifth immunization with IDLV expressing the CH505.w136 SOSIP gp140 Env, there was no boost in the antibody response induced by the previous immunization with non-stabilized gp140 Envs, suggesting that IDLV-CH505.w136 SOSIP immunization induced antibodies against epitopes only present on the closed envelope trimer. To confirm this, we performed PGT151 SOSIP capture ELISAs to detect anti-CH505.w136 SOSIP specific antibodies. As shown in **figure 1d**, anti-CH505.w136 SOSIP specific antibodies were elicited in all animals between 2 and 6 weeks post-vaccination and were significantly boosted (p = 0.0156 for both comparisons using Wilcoxon signed-rank tests) by a second IDLV-CH505.w136 SOSIP ^+/-^ protein immunization. These data suggest that immunization with IDLV-CH505.w136 SOSIP Env induced a narrower and more specific Ab response compared to immunizations with non-stabilized gp140 Envs.

### Epitope mapping of antibody response induced by sequential IDLV-CH505 immunizations

We next characterized the epitope specificities of the antibodies induced by the sequential IDLV-CH505 Envs ^+/-^ protein immunizations by peptide microarray. Serum samples collected at 2 weeks after each boost immunization (weeks 26, 51, 75, 97 and 113) were tested for their ability to bind to a panel of Env sequences from different clades. Immunized animals in both groups developed cross-clade responses against a range of linear epitopes in gp120 (**figure 2a**) and gp41 (**figure 2b**), including C1, V3, V5, C5, AVERY, gp41 IDR, and MPER. Binding against the variable loop 2 (V2) was also detected in all the animals but was primarily specific for CH505 sequences (**figure 2a**). Binding specificities were overall similar between the two groups (**supplementary figure 1**), but the magnitude of the responses trended higher in group B2 animals for the majority of the epitopes (**figure 2**). Following IDLV-CH505.w136 SOSIP boosts a reduction in the magnitude of binding responses against different epitopes was observed in both groups of animals (**figure 3**). Differences were observed for V2 and MPER binding responses between group B1 and B2 animals after the last IDLV-CH505.w136 SOSIP immunization (**figure 3b**). These data further suggest that immunization with IDLV-CH505.w136 SOSIP Env induced a narrower Ab response compared to immunizations with non-stabilized gp140 Envs and that the addition of the CH505.w136 SOSIP Env protein in group B2 animals resulted in higher magnitude of antibody binding against specific epitopes, including V2 and MPER.

**Figure 2.**
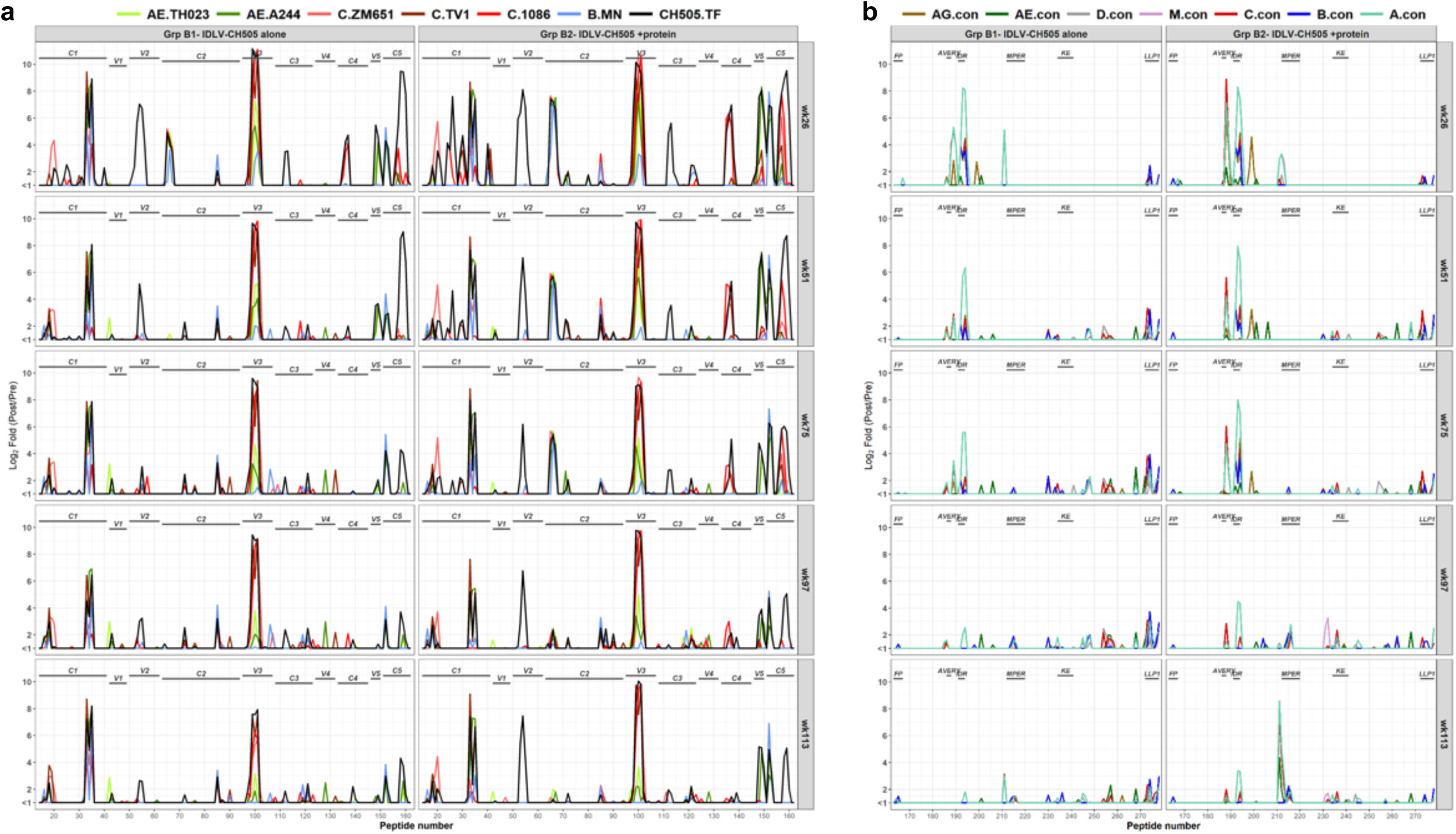
Linear epitope specificities of antibody responses induced by sequential IDLV-CH505 Env immunizations. gp120 (**a**) and gp41 (**b**) binding plots for serum samples from macaques immunized with either IDLV-CH505 alone or IDLV-CH505 + protein at 2 weeks after each immunization. Numbers on the x axis are peptide numbers in the array library. Different colors of bars represent different strains/clades as indicated in the keys in the panels (TH023, A244, ZM651, TV-1, 1086C, MN, CH505 for panel a and AG, AE, D, M, C, B and A consensus for panel b).

**Figure 3.**
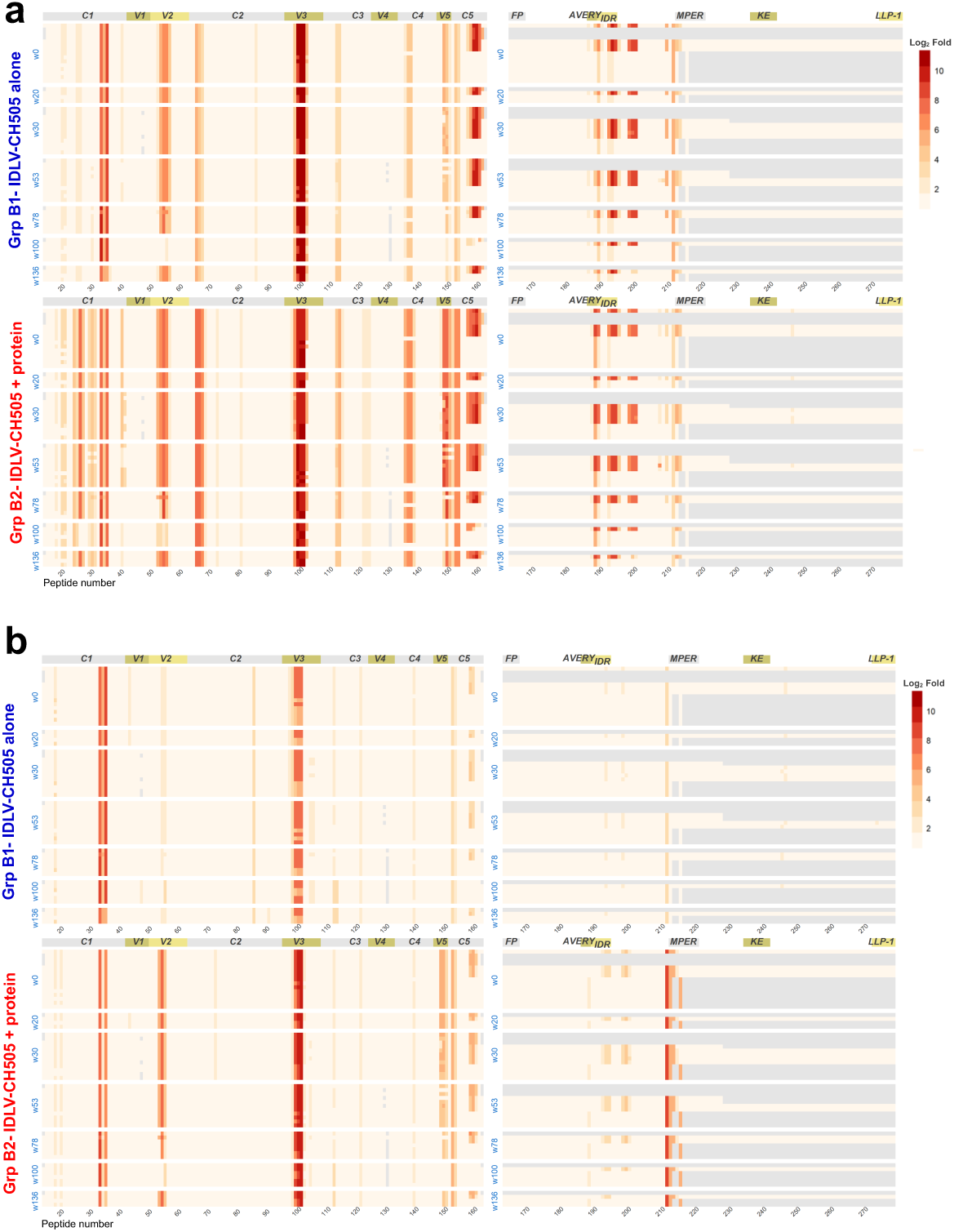
Linear epitope binding to CH505 sequences. The heat maps show gp120 and gp41 binding, at week 26 (**a**) and week 113 (**b**) post IDLV ^+/-^ protein immunization, to the different CH505 strain sequences used in immunization. Binding intensity is shown for each peptide, corrected with its own background value. Light grey shaded areas indicate sequence not present in the array library.

### Sequential IDLV-CH505 immunizations induce high-titers of tier-1 neutralizing antibody responses but no breadth

We next assessed the serum neutralizing activity induced by sequential IDLV-CH505 Envs immunization. Neutralization assays were performed on serum samples collected at different time points post-immunization including at the peak of the antibody response (weeks 6, 26, 51, 75, 101 and 113), 6 months after the second (week 49) and fourth (week 91) immunizations as well as 2, 11 and 16 weeks (weeks 97, 106 and 111 respectively) after the first IDLV-CH505.w136 SOSIP immunization and 2 weeks (week 113) after the second IDLV-CH505.w136 SOSIP immunization. Serum samples from all 16 macaques in groups B1 and B2 neutralized the autologous tier 1 virus CH505 w4.3 after the second immunization with IDLV-CH505.w53 gp140 Env immunization (**Table 1**). However, nAbs against the four sequential autologous viruses (CH505 TF, week 53, week 78, week 100) and the challenge SHIV CH505.375H were not elicited (data not shown). We also assayed sera starting at week 75 for neutralization of viruses that are sensitive to neutralization by the inferred unmutated common ancestor and intermediate antibodies in the CH103 lineage^21^, but failed to detect neutralization of this panel of viruses. Significantly higher tier 1 neutralizing antibody titers were detected in group B2 animals (IDLV-CH505 + protein) compared to group B1 animals (IDLV-CH505 alone) across all time points (p < 0.0001 using an aligned rank test). An increase in nAb titers was detected after each immunization except after the last IDLV-CH505.w136 SOSIP boost. These data indicate that sequential IDLV-CH505 Env immunizations induced high titer of tier 1 neutralizing antibodies but no neutralization breadth.

**Table 1.**
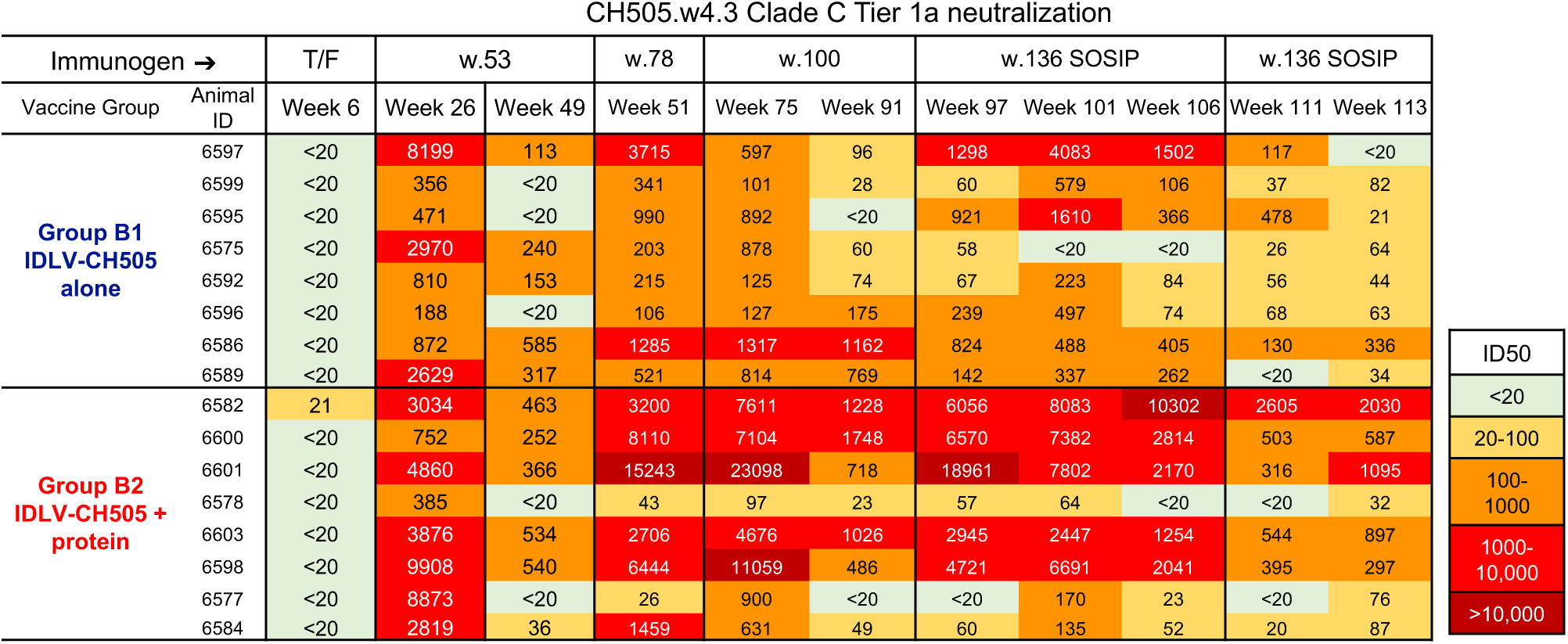
Serum neutralization activity against the clade C tier1 virus CH505.w4.3. Values are the serum dilutions at which relative luminescence units (RLU) were reduced by 50% compared to RLU in virus control wells after subtraction of background RLU in cell control wells. A response was considered positive if the post-immunization ID50 was 3 times higher than the pre-immune ID50 and 3 times greater than the signal against the MLV-pseudotyped negative control virus. No neutralization against other CH505 tier 1 and tier 2 viruses (CH505.T/F (tier 2), CH505w53.16 (tier 1a), CH505w.78 (tier 1b) and CH505w.100 (tier 2)) was detected.

CH505.w4.3 Clade C Tier 1a neutralization
| Immunogen → |  | T/F | w.53 |  | w.78 | w.100 |  | w.136 SOSIP |  |  | w.136 SOSIP |  |
| --- | --- | --- | --- | --- | --- | --- | --- | --- | --- | --- | --- | --- |
| Vaccine Group | Animal ID | Week 6 | Week 26 | Week 49 | Week 51 | Week 75 | Week 91 | Week 97 | Week 101 | Week 106 | Week 111 | Week 113 |
| **Group B1**<br>**IDLV-CH505**<br>**alone** | 6597 | <20 | 8199 | 113 | 3715 | 597 | 96 | 1298 | 4083 | 1502 | 117 | <20 |
|  | 6599 | <20 | 356 | <20 | 341 | 101 | 28 | 60 | 579 | 106 | 37 | 82 |
|  | 6595 | <20 | 471 | <20 | 990 | 892 | <20 | 921 | 1610 | 366 | 478 | 21 |
|  | 6575 | <20 | 2970 | 240 | 203 | 878 | 60 | 58 | <20 | <20 | 26 | 64 |
|  | 6592 | <20 | 810 | 153 | 215 | 125 | 74 | 67 | 223 | 84 | 56 | 44 |
|  | 6596 | <20 | 188 | <20 | 106 | 127 | 175 | 239 | 497 | 74 | 68 | 63 |
|  | 6586 | <20 | 872 | 585 | 1285 | 1317 | 1162 | 824 | 488 | 405 | 130 | 336 |
|  | 6589 | <20 | 2629 | 317 | 521 | 814 | 769 | 142 | 337 | 262 | <20 | 34 |
| **Group B2**<br>**IDLV-CH505 +**<br>**protein** | 6582 | 21 | 3034 | 463 | 3200 | 7611 | 1228 | 6056 | 8083 | 10302 | 2605 | 2030 |
|  | 6600 | <20 | 752 | 252 | 8110 | 7104 | 1748 | 6570 | 7382 | 2814 | 503 | 587 |
|  | 6601 | <20 | 4860 | 366 | 15243 | 23098 | 718 | 18961 | 7802 | 2170 | 316 | 1095 |
|  | 6578 | <20 | 385 | <20 | 43 | 97 | 23 | 57 | 64 | <20 | <20 | 32 |
|  | 6603 | <20 | 3876 | 534 | 2706 | 4676 | 1026 | 2945 | 2447 | 1254 | 544 | 897 |
|  | 6598 | <20 | 9908 | 540 | 6444 | 11059 | 486 | 4721 | 6691 | 2041 | 395 | 297 |
|  | 6577 | <20 | 8873 | <20 | 26 | 900 | <20 | <20 | 170 | 23 | <20 | 76 |
|  | 6584 | <20 | 2819 | 36 | 1459 | 631 | 49 | 60 | 135 | 52 | 20 | 87 |

**Table 1. Serum neutralization activity against the clade C tier1 virus CH505.w4.3.**
| ID50 |
| --- |
| <20 |
| 20-100 |
| 100-1000 |
| 1000-10,000 |
| >10,000 |

### Sequential IDLV-CH505 immunizations induce higher binding and neutralizing antibody titers than protein and DNA^+/-^ protein immunization regimens

We next compared the magnitude and durability of binding and neutralizing antibody responses between sequential IDLV ^+/-^ protein, protein alone and DNA ^+/-^ protein immunization regimens delivering the same sequential CH505 envelopes^22, 23^. The same adjuvant (GLA-SE) was used in all the regimens that included protein. The immunization schedule for each of the vaccine regimens tested is shown in **figure 4a.** Of note, while in the IDLV-based regimens the animals were immunized every 6 months, in the protein and DNA ^+/-^ protein regimens the animals were immunized every 6 weeks before receiving a delayed boost at either 4 or 8 months, respectively. Significantly higher magnitude of antibody responses were induced in the IDLV ^+/-^ protein groups compared to DNA ^+/-^ protein groups (**figure 4b**). The durability of antibody responses at six months post-vaccination was also significantly higher in the IDLV ^+/-^ protein groups compared to protein or DNA alone (4 months). Similarly, neutralizing antibody titers were significantly higher in the IDLV^+/-^ protein groups compared to DNA alone and in the IDLV + protein group compared to protein (**figure 4c**). Although the median nAb titer values for the IDVL + protein group was 5890 and for the DNA + protein group was 640, this difference did not reach statistical significance due to the small sample size of the DNA + protein group (n= 4). These comparisons demonstrate that IDLV^+/-^ protein induced a more durable antibody response compared to protein alone and DNA ^+/-^ protein vaccine regimens.

**Figure 4.**
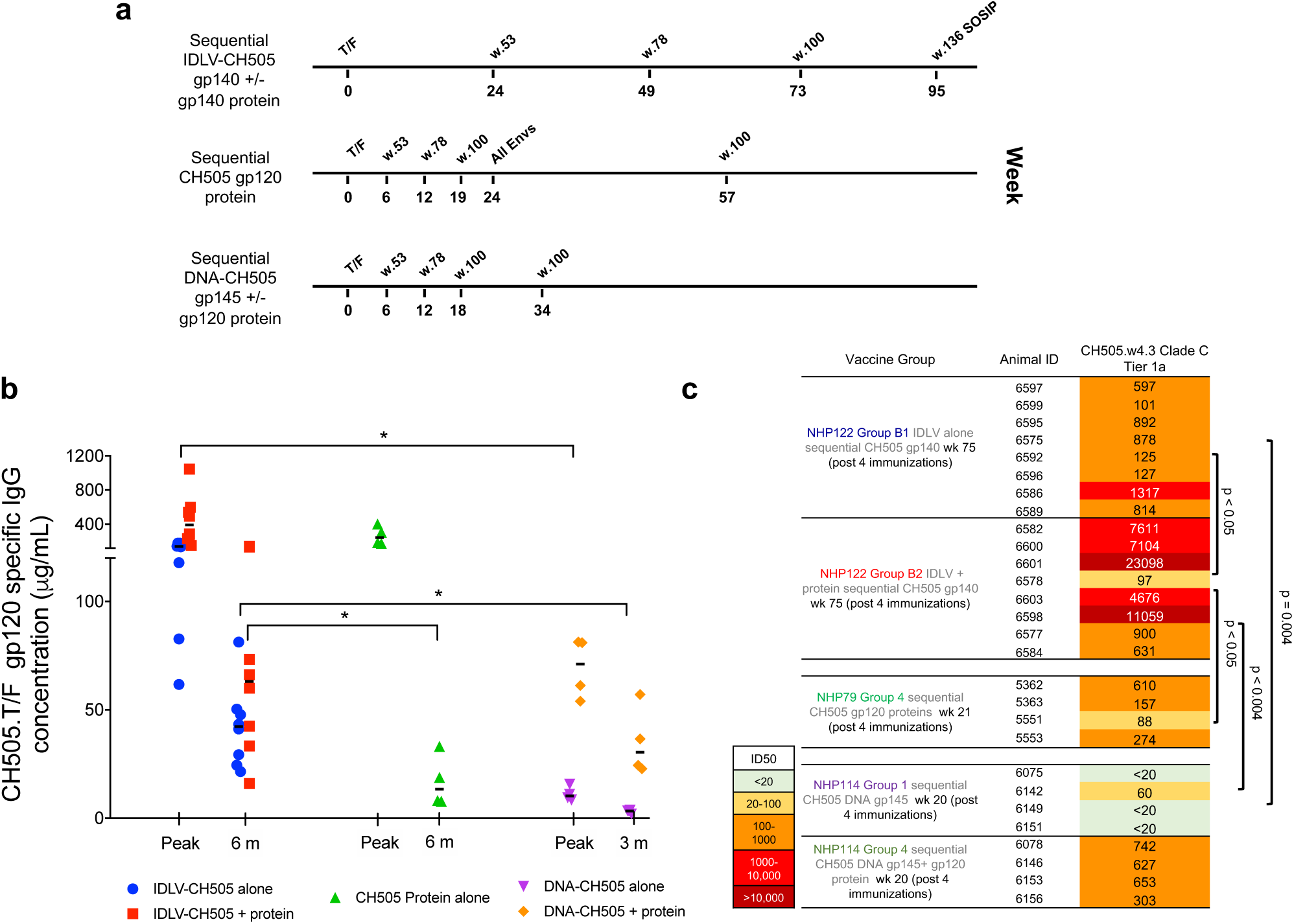
Magnitude and durability of antibody responses induced by sequential IDLV, protein and DNA ^+/-^ protein immunizations. **(a)** Non-human primate immunization regimens with sequential IDLV ^+/-^ protein, protein alone and DNA ^+/-^ protein. (**b**) ELISA binding of plasma antibodies to CH505 T/F Env at the peak and either six or three months (for DNA ^+/-^ protein) post four immunizations with each vaccine regimen. Binding titers measured as concentration in μg/mL starting at a 1:3000 plasma dilution. Asterisks indicate that a statistically significant difference was detected. (**c**) Serum neutralization activity against the clade C tier1 virus CH505.w4.3 post four immunizations with each vaccine regimen.

### Sequential IDLV-CH505 efficacy against autologous SHIV challenge

To assess the efficacy of the sequential IDLV ^+/-^ protein vaccines, we challenged animals in B1, B2 and B3 groups beginning at week 117 (6 weeks after the last immunization) ten times with a weekly intrarectal low-dose autologous Tier-2 SHIV.CH505.375H.dCT. Eleven animals immunized with an IDLV-GFP vector were challenged as the control arm. After ten challenges, only 1 of the 8 IDLV-CH505 + protein vaccinated animals remained uninfected, while all the animals in the IDLV-CH505 group became infected. In the control arm (IDLV-GFP) all the animals became infected after 8 challenges (**figure 5a**). There was a difference between IDLV-CH505 + protein, IDLV-CH505 alone and the IDLV-GFP control groups in peak viral load, with the IDLV-CH505 + protein group having ∼3.6 times lower peak viral load compared to IDLV-CH505 alone or IDLV-GFP (**figure 5b**). Furthermore, we observed more rapid and sustained control of viremia in the IDLV-CH505 + protein group compared to IDLV-CH505 alone group, with 71% (5/7) of group B2 animals and 37.5% (3/8) of group B1 animals having viral loads below the limit of detection at the time of necropsy (10 weeks post last challenge) (**figure 5c-d**). Virus control was observed also in 45.4% (5/11) of the animals in the control arm (IDLV-GFP) (**figure 5e**). As some of the vaccinated animals showed sustained viral load levels after infection, we analyzed the data from neutralization assays against the SHIV CH505.375H challenge virus at week 113 (prior to challenge) for enhancement of virus infection in vitro but detected none (data not shown).

**Figure 5.**
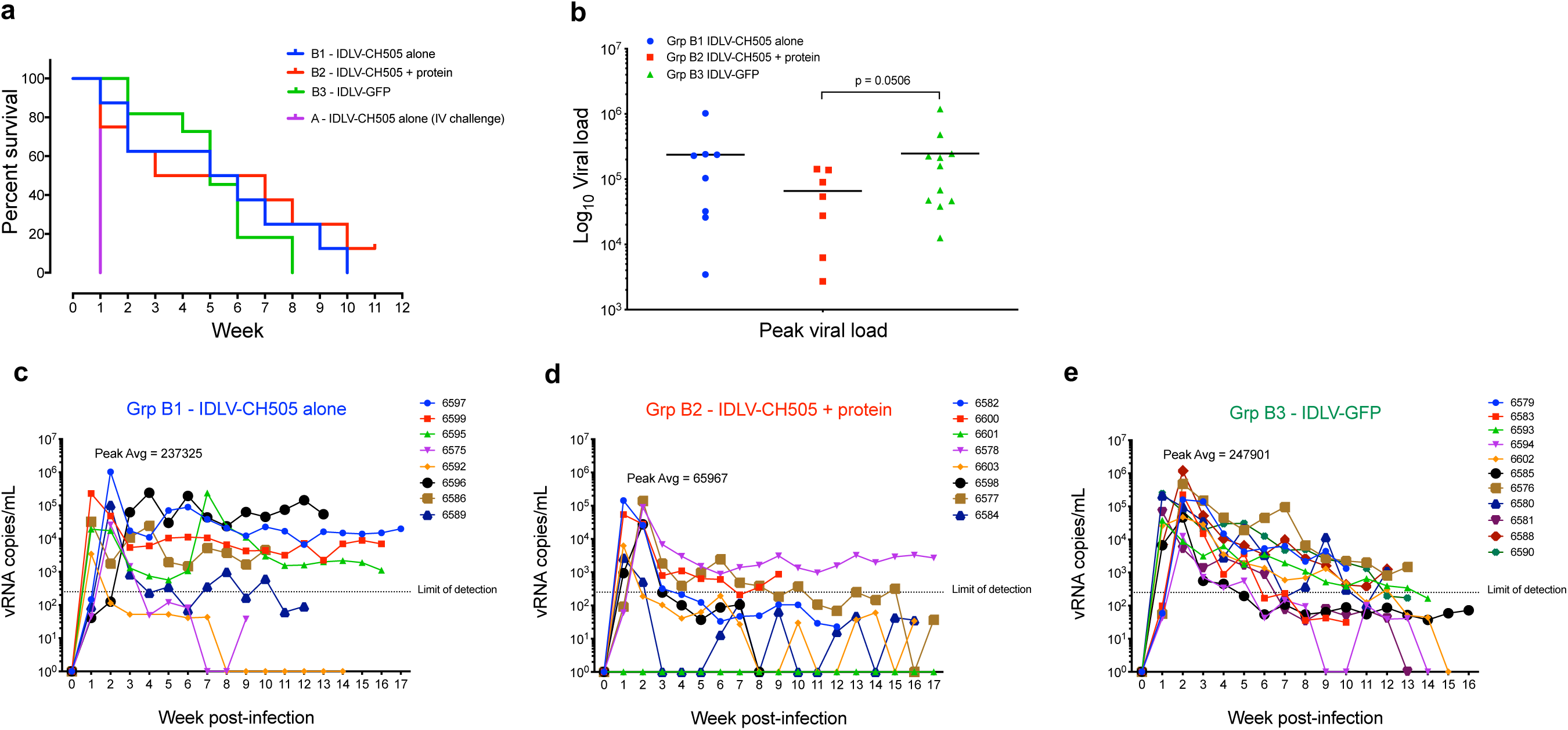
SHIV challenge outcomes in IDLV-CH505 ^+/-^ protein vaccinated animals. **(a)** Six weeks after completion of the scheduled vaccinations (week 117) the 16 macaques in the B1 and B2 vaccine groups and the 11 macaques in the B3 control group were challenged intrarectally with the Clade C tier 2 SHIV.CH505.375H.dCT. The four animals in group A were challenged once intravenously. The Kaplan-Meier plot shows the percentage of uninfected macaques after 10 weekly IR challenges in each group. (**b**) Peak viral load of the infected macaques from the vaccine and control groups. Lines are group means and (p=0.0506, Exact-Wilcoxon test). (**c-e**) Viral load trends from time to infection in each group. Each line represents one animal.

Because SIV-Gag is present in both the IDLV-CH505 and IDLV-GFP vector particles, we next measured anti SIV-Gag specific T cell responses in the three groups of animals, to determine whether differences in SIV-Gag responses between animals that controlled or did not control viral load could be detected. Similar anti SIV-Gag responses were detected in the three groups of animals (**supplementary figure 2 a-c**). The anti-CH505 Env T cell responses were also similar between group B1 (IDLV-CH505 alone) and group B2 (IDLV-CH505 + protein) animals (**supplementary figure 2 d-e**). No correlation between IFN-γ responses (weeks 75, 97 and 101) and time to virus acquisition was observed. These data suggest that the vaccine induced T cell responses did not influence the time to infection.

### Binding and neutralizing antibody titers correlate with time to virus acquisition

To determine whether the differences in binding and neutralizing antibody titers detected between the two vaccine groups played a role in the observed challenge outcome, we performed Kendall Tau correlation analyses. A correlation between binding antibody titers at weeks 75 and 97 (**figure 6a**) and time to virus acquisition (**figure 6b**) was observed (Kendall tau correlation p <0.05) for group B2 animals (IDLV-CH505 + protein). The animal that remained uninfected (6601) and the one that was infected at the last challenge (6600) demonstrated the higher magnitude of binding antibody responses (**figure 6a**), while the animals that got infected between 1 and 2 challenges (6577, 6578, 6584) had the lowest antibody titers. Similarly, a correlation between neutralizing antibody titers at weeks 101 for group B1 and weeks 75, 97 and 101 for group B2 (**figure 6c**) and time to virus acquisition was observed (Kendall tau correlation p <0.05).

**Figure 6.**
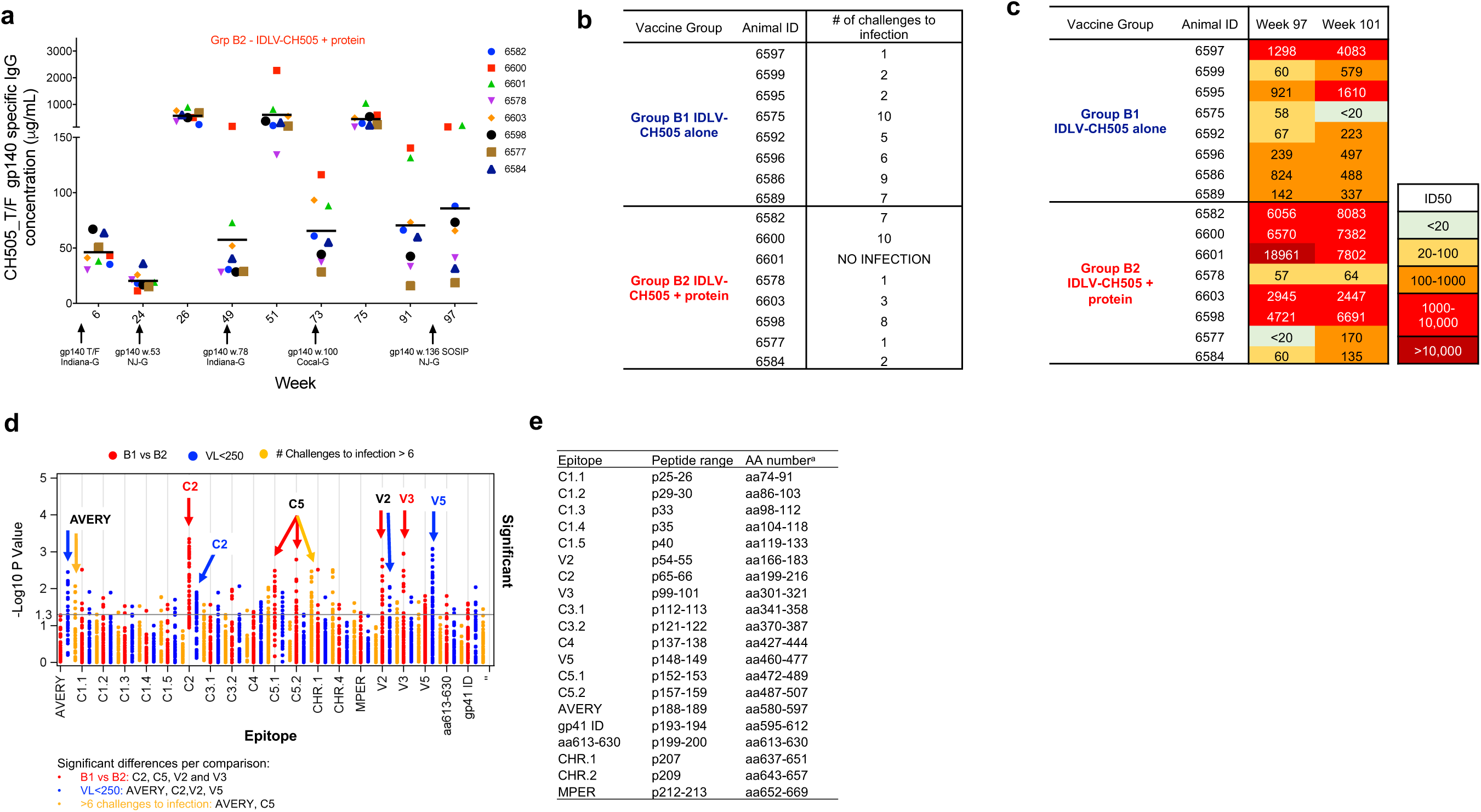
Magnitude and specificity of antibody responses in IDLV + protein vaccinated animals correlate with time to virus acquisition. **(a)** ELISA binding of plasma antibodies to CH505 T/F Env at the peak and six months post-each immunization with IDLV-CH505 + protein (group B2 animals). Each dot represents one animal (**b**) Number of challenges to virus acquisition for each animal in groups B1 and B2. (**c**) Serum neutralization activity against the clade C tier1 virus CH505.w4.3 at the indicated time points. (**d**) Comparison of linear epitope specificities between animals in B1 and B2 vaccine groups (red dots), animals with viral loads above or below 250 copies/mL (blue dots) and animals that required either less than 6 or more than 6 challenges to become infected (yellow dots). Significant differences fall above the horizontal line and are listed in the text box below the figure for each comparison. (**e**) The peptide range and the aa position for each epitope are indicated.

Because we observed quantitative differences in epitope binding specificities between the two vaccine regimens (**figure 3**), we next compared epitope binding responses among animals based on vaccine group (B1 vs B2), viral load (below and above 250 copies/mL) and time to acquisition (higher and lower than 6 challenges to infection). We observed significant differences (Exact Wilcoxon test) in epitope binding responses between group B1 and B2 for C2, C5, V2 and V3 epitopes (**figure 6d-e**). When comparing epitope binding specificities between controller vs non controller animals, significant differences in binding to AVERY, C2, V2 and V5 epitopes were observed. Additionally, the animals that required more than 6 challenges to infection demonstrated higher binding to AVERY and C5 epitopes.

These data suggest that the differences in nAb titers and magnitude of Ab specific for certain epitopes between the two vaccine regimens played a role in the challenge outcome.

### Absence of mobilization and or recombination between IDLV and the challenge virus

To evaluate the safety of sequential IDLV-CH505 Env immunization a group of 4 macaques was challenged intravenously with SHIV.CH505.375H.dCT (**figure 1a**) to enhance the possibility of potential recombination and/or mobilization events between the challenge virus and the IDLV-CH505 vaccines. All the animals became infected after one IV challenge (**figure 5a**). Mean peak plasma viral load for the group was 136,322 copies/mL and at the time of necropsy (12 weeks post-challenge) 2 of the 4 animals had viral loads below 250 copies/mL (**figure 7a**). To assess the possibility of recombination and/or mobilization events, we performed single genome amplification (SGA) on peripheral blood mononuclear cells (PBMC) and lymph node cells using primer sets designed to amplify any of the five CH505 gp140 *env* sequences encoded by IDLV (**supplementary figure 3**). We could not amplify any vector sequence in either PBMC or lymph node cells. Conversely, using SHIV.CH505.375H.dCT specific primers we were able to amplify several SHIV sequences in both PBMC and LN cells (**figure 7 b-c**). These data demonstrate no tendency for IDLV to recombine with actively replicating virus in an infected host and indicate that IDLV is a safe platform for HIV-1 vaccine that can be used in repeated immunizations.

**Figure 7.**
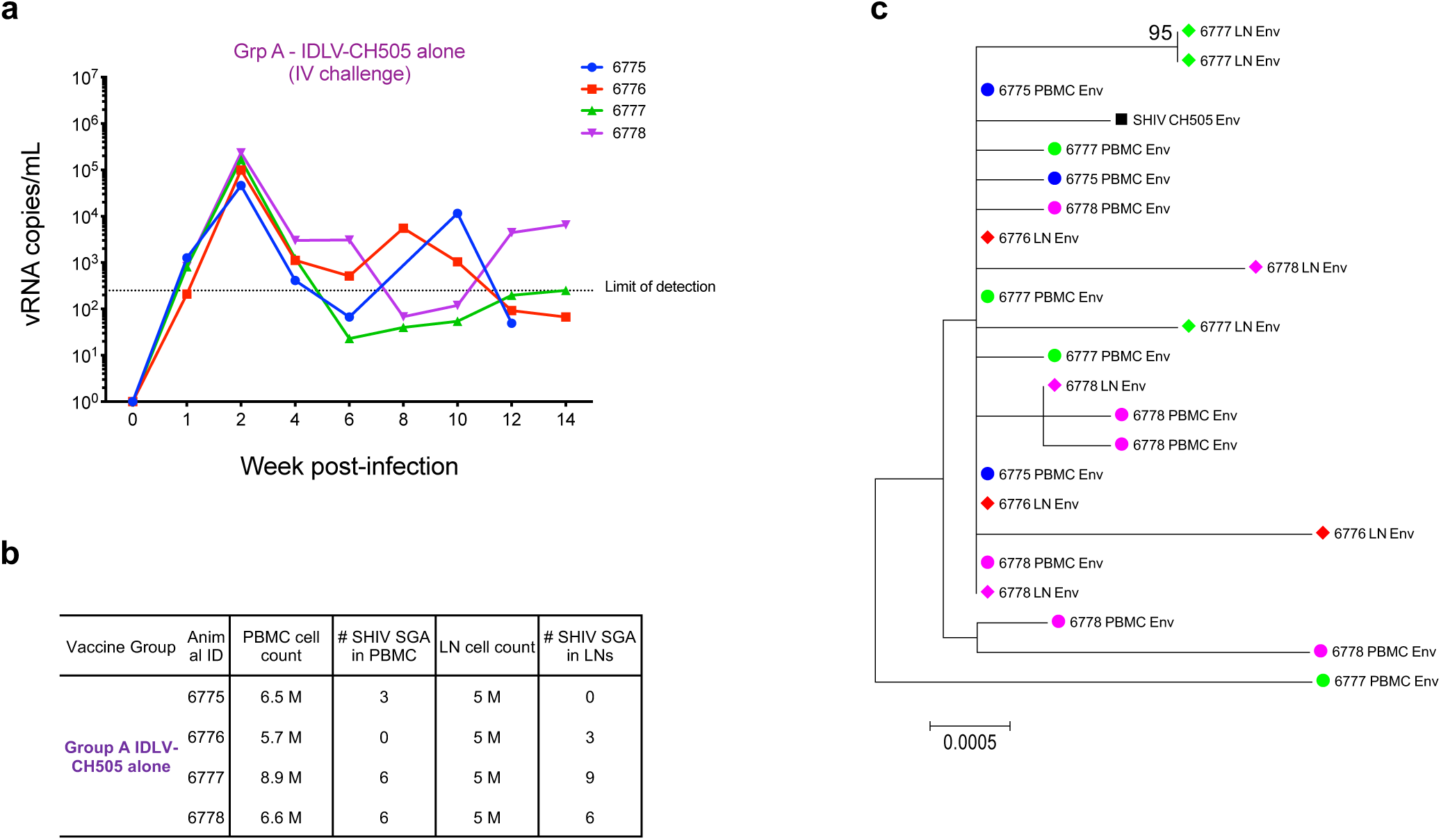
Absence of vector mobilization or recombination in IDLV-CH505 vaccinated animals challenged intravenously with SHIV-CH505. **(a)** Viral load trends from time of infection in group A animals (IDLV-CH505 alone). Each line represents one animal. (**b**) Number of SHIV.CH505.375H.dCT sequences amplified from either PBMC or lymph nodes samples for each animal. (**c**) Neighbor-joining phylogenetic tree including CH505 env sequences amplified from either PBMC (full circles) or lymph nodes (full diamonds) from each animal. Each color represents one animal. Bootstrap values ≥ 80 are shown. Genetic distance is indicated at the bottom of the figure and represents the number of nucleotide substitutions per site.

## Discussion

In this study, we evaluated the immunogenicity, safety and efficacy of sequential immunizations of rhesus macaques with an SIV-based IDLV, with or without protein, expressing a series of envelopes isolated from the CH505 individual who made the CH103 and CH235 broadly neutralizing antibody (bnAb) lineages. Our results show that co-immunization with IDLV and protein induced higher magnitude and durability of antibody responses compared to IDLV alone. In the IDLV + protein co-immunized animals we detected higher titers of neutralizing antibodies and higher V2 and MPER epitope binding after the last IDLV-CH505.w136 SOSIP immunization, suggesting that the addition of the protein improved neutralizing Abs titers and the magnitude of Abs against specific epitopes in the vaccinated macaques.

We also compared the magnitude and durability of the antibody response induced by IDLV-CH505 ^+/-^ protein vaccination to those induced by protein and DNA ^+/-^ protein vaccine regimens delivering the same series of CH505 envelopes in historical NHP studies ^22, 23^. Our analysis showed that co-immunization with IDLV and protein was superior to any of the tested vaccine regimens in terms of magnitude and durability of the immune responses at 6 months post-vaccination. Furthermore, IDLV alone induced higher magnitude of antibody responses compared to DNA ^+/-^ protein and more durable antibody responses compared to protein alone or DNA ^+/-^ protein. Similarly, serum neutralizing activity was significantly higher in the IDLV + protein co-immunized animals compared to all the other tested vaccine regimens. These data support the use of IDLV as a vaccine platform to induce higher magnitude and more durable immune responses against HIV-1 as well as other pathogens.

Consistent with previous studies that performed sequential immunizations of macaques with gp120 or gp140 CH505 envelopes^22, 23^, we did not detect neutralization breadth in the sera of the vaccinated animals in this study. Several studies have shown that SOSIP trimers engineered for increased Env stability better mimic native, virion-associated trimers both antigenically and structurally, including by displaying the epitopes for many bnAbs^4, 19, 20, 24, 25^. We therefore performed the last 2 immunizations with IDLV expressing a stabilized gp140 SOSIP Env trimer (CH505.w136 SOSIP) with the goal of focusing antibody responses towards the epitopes targeted by bnAbs. We observed that following IDLV-CH505.w136 SOSIP immunization there was induction of antibodies against epitopes only present on the closed envelope trimer, while we did not detect a boost in the antibody response induced by the previous immunization with non-stabilized gp140 Envs.

Although we detected a narrower and more specific Ab response compared to immunizations with non-stabilized gp140 Envs, the two immunizations with IDLV-CH505.w136 SOSIP Env were not sufficient at inducing neutralization breadth. Recent studies demonstrated that there are several other factors working against the elicitation of bnAbs by vaccination, including host control, the paucity of bnAb precursors, the structure of the HIV-1 Env and the need for bnAbs to acquire improbable somatic mutations for their neutralization activity^26-28^. Substantial progress has been made in immunogen design to overcome some of those roadblocks. Saunders *et al.* recently demonstrated that a structurally modified CH505 Env could bind to the initial precursor of the bnAb lineage and could select for specific HIV-1 bnAb improbable mutations, resulting in the induction of potent neutralizing CD4 binding site antibodies in macaques^29^. Future studies will determine whether the expression of these novel immunogens by IDLV will drive bnAb maturation.

Although the induction of bnAbs by vaccination is desirable, results from human HIV-1 vaccine trials (RV144, APPROACH and IPCAVD010/HPX1002) as well as from preclinical studies in NHPs suggest that non-neutralizing antibodies could also reduce the risk of virus acquisition and mediate protection^3, 30-32^. We therefore evaluated the efficacy of sequential IDLV-CH505 ^+/-^ protein vaccination against autologous virus acquisition. Only one animal in the IDLV-CH505 + protein co-immunization group resisted infection following 10 intrarectal challenges with SHIV.CH505.375H.dCT. This group of animals demonstrated 3.6 times lower peak viremia and better virus control compared to the control group of animals immunized with IDLV-GFP or IDLV-CH505 alone. Interestingly, we observed a positive correlation between the magnitude of binding and neutralizing antibody responses and time to virus acquisition. Significant differences in binding to AVERY, C2, V2 and V5 epitopes were also observed between controller and non-controller animals. Additionally, animals that required more than 6 challenges to become infected demonstrated stronger binding to AVERY and C5 epitopes. All together these data suggest that the differences in the magnitude of the antibody responses induced by the two vaccine regimens (IDLV-CH505 alone and IDLV-CH505 + protein) resulted in a more favourable challenge outcome in the co-immunization group.

Another important aspect we addressed in this study is the safety of the IDLV-vaccine following infection with a replication competent virus. While there are a number of safety features built into the SIV-based IDLV used for immunization, testing these features in the setting of viral infection is critical, especially regarding the chance for a recombination event resulting in replication competent virus. We have previously demonstrated that immunization of rhesus macaques with IDLV-Env did not result in the generation of RCLs in vivo^11^. Here, we assessed the safety of sequential IDLV vaccinations by evaluating vector mobilization and/or recombination events in PBMC and lymph nodes samples of 4 macaques vaccinated with sequential IDLV-CH505 and then challenged intravenously with SHIV.CH505. No evidence of vector mobilization or recombination between IDLV and the challenge virus could be detected.

In summary, the use of IDLV as a vaccine delivery system to express HIV-1 Env is a promising approach, particularly in light of its safety profile, the higher durability of immune responses, higher magnitude of neutralization and persistence of antigen expression compared to DNA and protein vaccination strategies. While IDLV proved successful at inducing high titer and durable immune responses, the immunogen strategy utilizing sequential non stabilized/optimized trimers did not induce tier 2 antibody responses, and this has been consistent across the different vaccine platforms used to deliver sequential gp120 or gp140 CH505 Envs. However, recent data from our group and others have demonstrated that conformationally stabilized Env trimer immunogens, in contrast to non-stabilized Envs, consistently induce autologous tier-2 neutralizing antibodies^4, 29, 33^. Future studies will evaluate the ability of IDLV delivering stabilized Env trimers to enhance B-cell maturation and drive bnAb development.

## Methods

### Construction of IDLV-CH505 Envs plasmids

The clade C HIV-1 Env C.CH505 gp140 glycoproteins (T/F, w.53, w.78. w.100 and w.136 SOSIP)^5^ were cloned into a SIV-based self-inactivating lentiviral transfer vector downstream of the internal CMV promoter (pGAE-CMV-CH505-Env-Wpre). The transfer vector pGAE-CMV-GFP-Wpre expressing the green fluorescent protein (GFP), the IN-defective packaging plasmids pAd-SIV-D64V, and the vesicular stomatitis virus envelope glycoproteins (VSV-G) for vector pseudotyping have been previously described^12, 34^.

### Vector production and validation

The human epithelium kidney 293T Lenti-X cells (Clontech Laboratories, Mountain View, CA) were maintained in Dulbecco’s Modified Eagles medium (Thermo Fisher Scientific, Waltham, MA) supplemented with 10% fetal bovine serum (GE Healthcare Life Sciences, HyClone Laboratories, South Logan, UT) and 100 units per ml of penicillin–streptomycin–glutamine (PSG) (Thermo Fisher Scientific). For production of recombinant IDLV, 3.5 × 10^6^ Lenti-X cells were seeded on 100 mm diameter Petri dishes and transfected with 12 µg per plate of a plasmid mixture containing transfer vector, packaging plasmid and VSV.G plasmid in a 6:4:2 ratio, using the JetPrime transfection kit (Polyplus Transfection Illkirch, France) following the manufacture’s recommendations. At 48 and 72 hours post-transfection, culture supernatants were cleared from cellular debris by low-speed centrifugation and passed through a 0.45 μm pore size filter unit (Millipore, Billerica, MA). Filtered supernatants were concentrated by ultracentrifugation for 2 hours at 23.000 RPM on a 20% sucrose cushion. Pelleted vector particles were resuspended in 1× phosphate-buffered saline (PBS) and stored at −80 °C until further use. Each IDLV stock was titered using a reverse transcriptase (RT) activity assay (RetroSys RT ELISA kit, Innovagen, Lund, Sweden) and the corresponding transducing units (TU) calculated by comparing the RT activity of each IDLV-CH505-Envs stock to the RT activity of IDLV-GFP stocks with known infectious titers^11^.

### Animals, IDLV-injection and SHIV challenge protocol

The thirty-one Indian origin rhesus macaques (*Macaca mulatta*) used in this study were negative for *Mamu-A*01, Mamu-B*08* and *Mamu-B*17* alleles and were housed at BIOQUAL, Inc. in accordance with the recommendations of the Association for Assessment and Accreditation of Laboratory Animal Care International Standards and with the recommendations in the Guide for the Care and Use of Laboratory Animals of the United States - National Institutes of Health. The Institutional Animal Use and Care Committee of BIOQUAL approved these experiments (study # 18-001).

Animals were immunized six times intramuscularly with sequential IDLV-CH505-Envs with 3 × 10^8^ TU per animal in 0.7-ml injection volume divided into two sites (left and right quadriceps). In the IDLV-CH505 + protein group the animals received 100 μg of the corresponding CH505 gp140 protein in 25 μg of GLA-SE adjuvant (synthetic monophosphoryl lipid A in stable emulsion)^18^ in addition to IDLV. Anticoagulated peripheral blood was obtained prior to IDLVs injection, 2 weeks post each immunization and monthly thereafter. Animals in groups B1, B2 and B3 were challenged ten times weekly by the intrarectal route with 1:100 dilution of SHIV.CH505.375H.dCT^16^ challenge stock. Animals in group A were challenged once intravenously with the same SHIV.CH505.375H.dCT dose. The virus stock was grown from the infectious molecular clone in rhesus peripheral blood mononuclear cells (PBMCs) and the stock was titrated in rhesus macaques to select the appropriate dilution.

SHIV plasma viral RNA measurements were performed at the Immunology Virology Quality Assessment Center Laboratory Shared Resource, Duke Human Vaccine Institute, Durham, NC as described.

### Direct ELISAs

High binding EIA/RIA 384 well plates (Corning) were coated overnight with 2 µg ml^-1^ of CH505.T/F gp140 protein in coating buffer (KPL, Gaithersburg, MD). After 1 wash with washing buffer (1X PBS + 0.1% Tween 20) plates were treated for 1 hour at room temperature with 40 μl per well of blocking buffer (PBS containing 4% [wt/vol] whey protein–15% normal goat serum–0.5% Tween 20). Serial 3-fold dilutions of plasma (from 1:3000 to 1:729000) or monoclonal antibodies (from 100 μg mL^-1^ to 0.5 ng mL^-1^) in blocking buffer were added to the plates (10 μl per well) in duplicates and incubated for 1.5 hours at room temperature. The Rhesus B12 IgG (b12R1) was used to develop standard curves (range 100 to 0.005 ng mL^-1^, with each dilution assayed in duplicate). Abs were detected by adding 10 μl per well of horse radish peroxidase (HRP)-conjugated, polyclonal goat anti-monkey IgG (Rockland, Gilbertsville, PA) diluted in blocking buffer (1:6000) and by adding 20 μl per well the SureBlue Reserve TMB microwell peroxidase substrate and stop solution (KPL, Gaithersburg, MD). Binding titers were analyzed as area-under-curve of the log transformed concentrations (Softmax Pro 7, Molecular Devices LLC, CA).

For detection of CH505.w136 SOSIP gp140 specific antibodies in monkeys’ sera, PGT151 SOSIP capture ELISAs were performed as described^29^.

### Linear epitope mapping microarray

Specificity and magnitude of binding response to cross-subtype linear epitopes were measured in a peptide microarray assay as previously described^35^ with minor modifications. The library contains overlapping peptides (15-mers overlapping by 12) covering 7 full length gp160 consensus sequences (subtype A, B, C, D, Group M, CRF1, and CRF2 consensuses), gp120 sequences of 6 vaccine strains (MN, A244, Th023, TV-1, ZM641, 1086C), and sequential CH505 Env strains as gp120, gp140, and SOSIP sequences. Microarray scan images were analyzed using MagPix 8.0 software to obtain binding intensity values for all peptides. Magnitude of binding is calculated as the log2 fold difference, post-/pre-immunization intensity.

### HIV Neutralization assays

Neutralization of Env-pseudotyped viruses was measured in 96-well culture plates using Tat-regulated firefly luciferase (Luc) reporter gene expression to quantify reductions in virus infection in TZM-bl cells^36^. Serum neutralization was measured against: SVA-MLV (negative control for non-specific activity in the assay), CH505.w4.3 (tier 1), CH505.T/F (tier 2), CH505w53.16 (tier 2), CH505w.78 (tier 1b) and CH505w.100 (tier 2); SHIV CH505.375H (tier 2), CH505TF.gly3.276/GnTI- (tier 2), CH505TF.gly4/293S-GnTI- (tier 1b) and CH505s.G458Y.N279K/293S-GnTI- (tier 2). Heat-inactivated (56 °C, 1 hour) serum samples were assayed at threefold dilutions starting at 1:20. Neutralization titers (50% inhibitory dose (ID50)) are the serum dilutions at which relative luminescence units (RLU) were reduced by 50% compared to RLU in virus control wells after subtraction of background RLU in cell control wells. A response was considered positive if the titer of the post-immunization ID50 was 3 times higher than that of the pre-immune ID50 and 3 times greater than the signal against the MLV-pseudotyped negative control virus.

### IFN-γ ELISpot assay

Multiscreen ninety-six well plates were coated overnight with 100 μl per well of 5 μg/ml anti-human interferon-γ (IFN-γ) (B27; Becton, Dickinson and Company, Franklin Lakes, NJ) in endotoxin-free Dulbecco’s-PBS (D-PBS). Plates were washed three times with D-PBS containing 0.1% Tween-20, blocked for 2 hours with Roswell Park Memorial Institute 1640 medium (RPMI) containing 10% fetal bovine serum and incubated with peptide pools and 2 × 10^5^ PBMCs in triplicate in 100 μl reaction volumes. Each peptide pool (1 μg/ml) was comprised of 15 amino acid peptides overlapping by 11 amino acids. The pools covered the entire HIV-Env and SIV-Gag proteins. Following 18-hour incubation at 37 °C, plates were washed nine times with Dulbecco’s PBS containing 0.1% Tween-20 and once with distilled water. Plates were then incubated with 2 μg/ml biotinylated rabbit anti-human IFN-γ (U-CyTech biosciences, Utrecht, The Netherlands) for 2 hours at room temperature, washed six times with D-PBS containing 0.1% Tween-20, and incubated for 1.5 hours with a 1:500 dilution of streptavidin-AP (Southern Biotechnology, Birmingham, AL). After five washes with D-PBS containing 0.1% Tween-20 and three washes with D-PBS alone, the plates were developed with bromochloroindolyl phosphate–nitro blue tetra-zolium (BCIP-NBT) chromogen (Thermo Fisher Scientific), stopped by washing with tap water, air dried, and read with an ELISpot reader (Cellular Technology Limited (CTL), Cleveland, OH) using ImmunoSpotAnalyzer software (Cellular Technology Limited (CTL)). Samples were considered positive if number of spot forming cells (SFC) was above twice that of the background (unstimulated) and > 50 SFC per million PBMC.

### Vector and SHIV detection in PBMC and LNs

DNA was isolated from PBMC and lymph nodes cells collected 12 weeks post-challenge using the QIAamp micro kit (Qiagen) following the manufacturer’s instructions and eluted in 50 μL of water. Standard curves for LV-CH505 DNA were generated using serial dilutions of genomic DNA extracted from 293T cells transduced with an integrase competent lentiviral vector expressing the CH505 T/F gp140 envelope (293T-LV-CH505). PCR reactions were performed using the primer sets and conditions shown in **supplementary figure 2**. All alignments were made using gene cutter (hiv.lanl.gov) and phylogenetic trees were made with MEGA6^37^. Neighbor-joining trees were constructed using the Kimura 2-parameter mode and the reliability of topologies was estimated by performing bootstrap analysis with 1000 replicates.

### Statistical analysis

Comparisons between groups were made using exact Wilcoxon tests because of the small sample size. Comparisons within groups across time points were made using Wilcoxon signed-ranked tests. An aligned rank test was used to compare neutralizing antibody titers across time points. Correlations were assessed using Kendall’s Tau correlation test. No adjustment to the alpha level was made for multiple comparisons. All computations were made using SAS v9.4 (SAS Institute, Inc.).

## Supplementary Figures

**Supplementary Figure 1.**
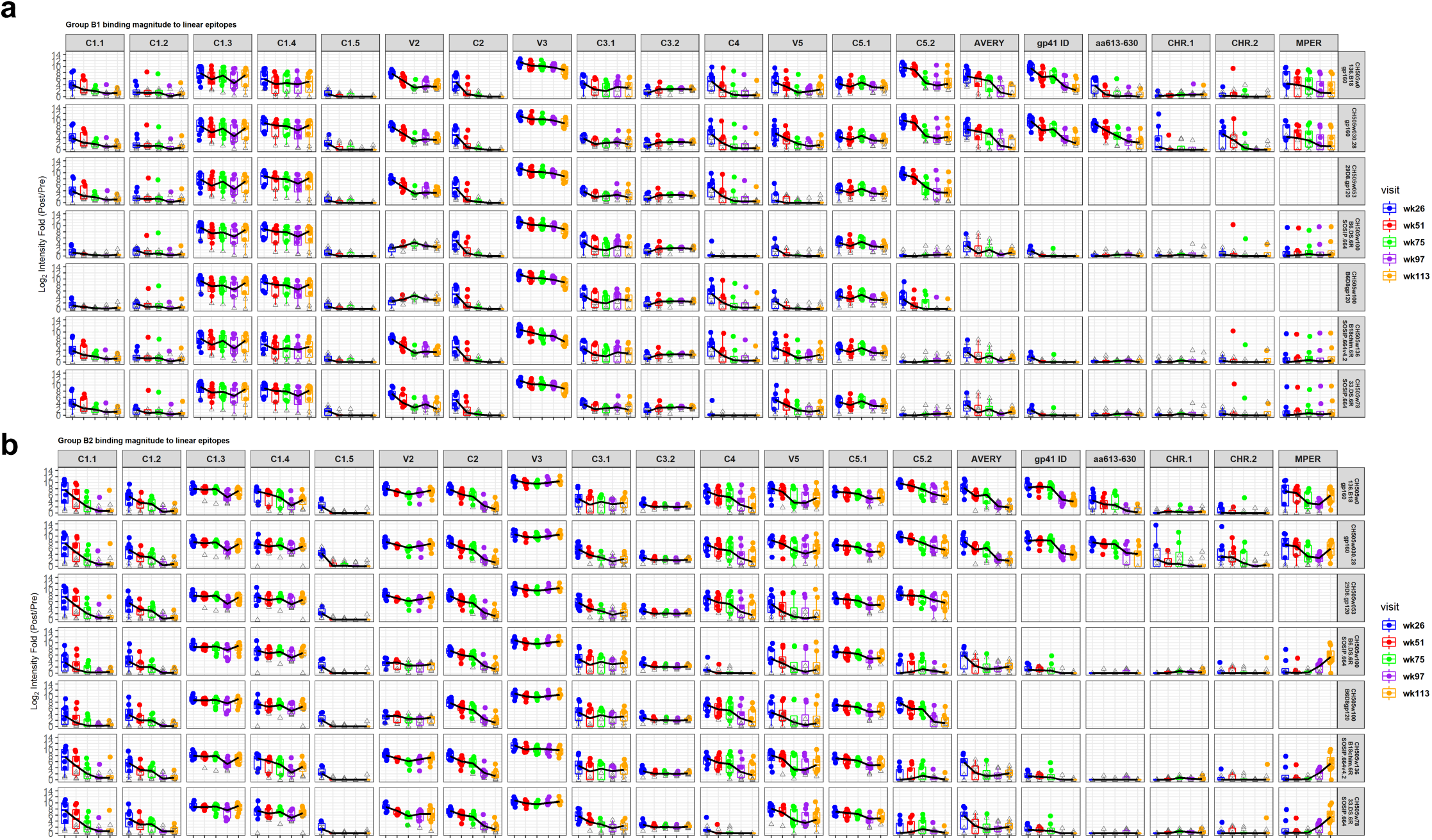
Linear epitope binding to CH505 sequences in IDLV- CH505 ^+/-^ protein sequentially vaccinated animals. Linear epitope specificities at 2 weeks after each immunization are shown for both IDLV-CH505 alone (**a**) and IDLV- CH505 + protein (**b**) immunized animals. Each color indicates a different time point. Binding intensity is shown for each peptide, corrected with its own background value.

**Supplementary Figure 2.**
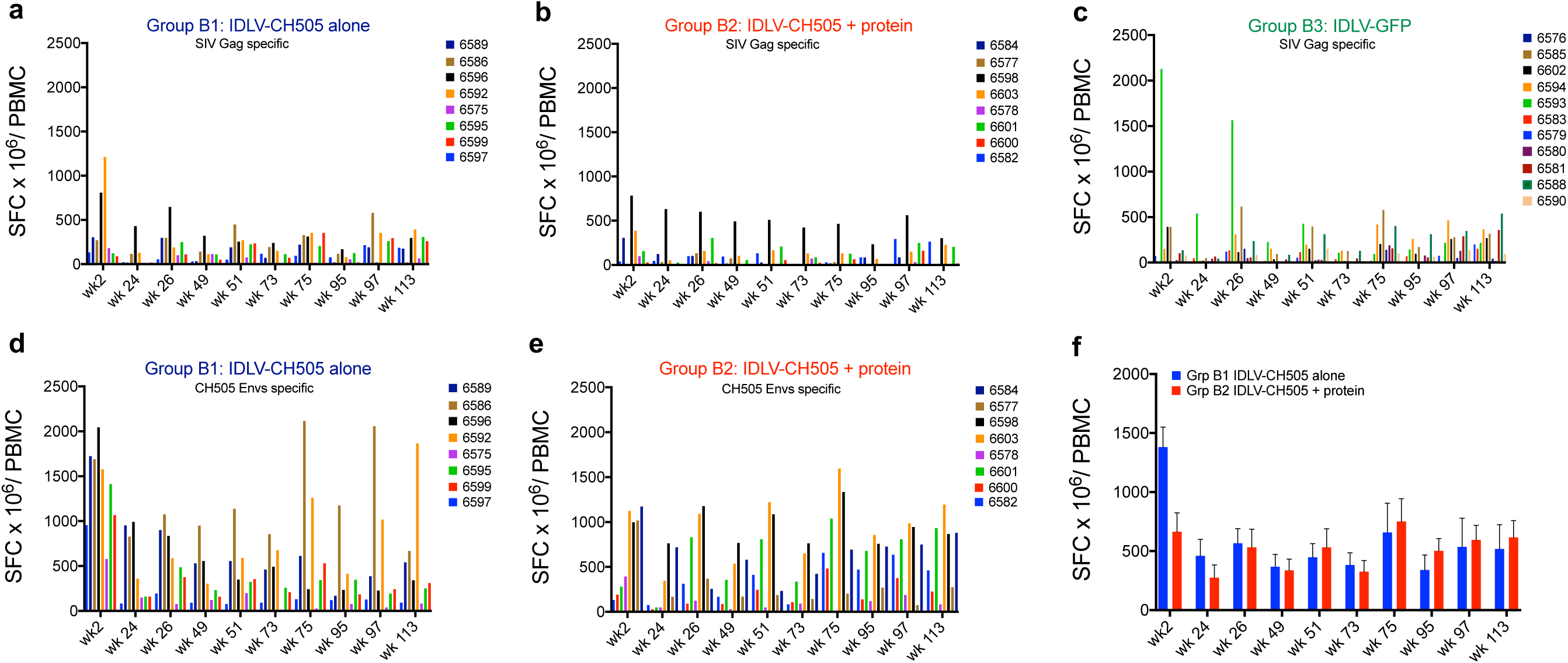
Env and Gag specific T-cell responses induced by IDLV- CH505 ^+/-^ protein and IDLV-GFP immunizations. **(a-c)** SIV-Gag and (**d-e**) CH505 Envs specific T cell responses were measured by IFN-γ ELISPOT in both the IDLV-CH505 ^+/-^ protein and IDLV-GFP vaccinated animals. Results are expressed as IFN-γ secreting cells, measured as spot forming cells (SFC) per million PBMC, at different time points post vaccination. Each line represents one animal. (**f**) Group mean IFN-γ ELISPOT anti-Env responses over time. Error bars indicate S.E.M.

**Supplementary Figure 3.**
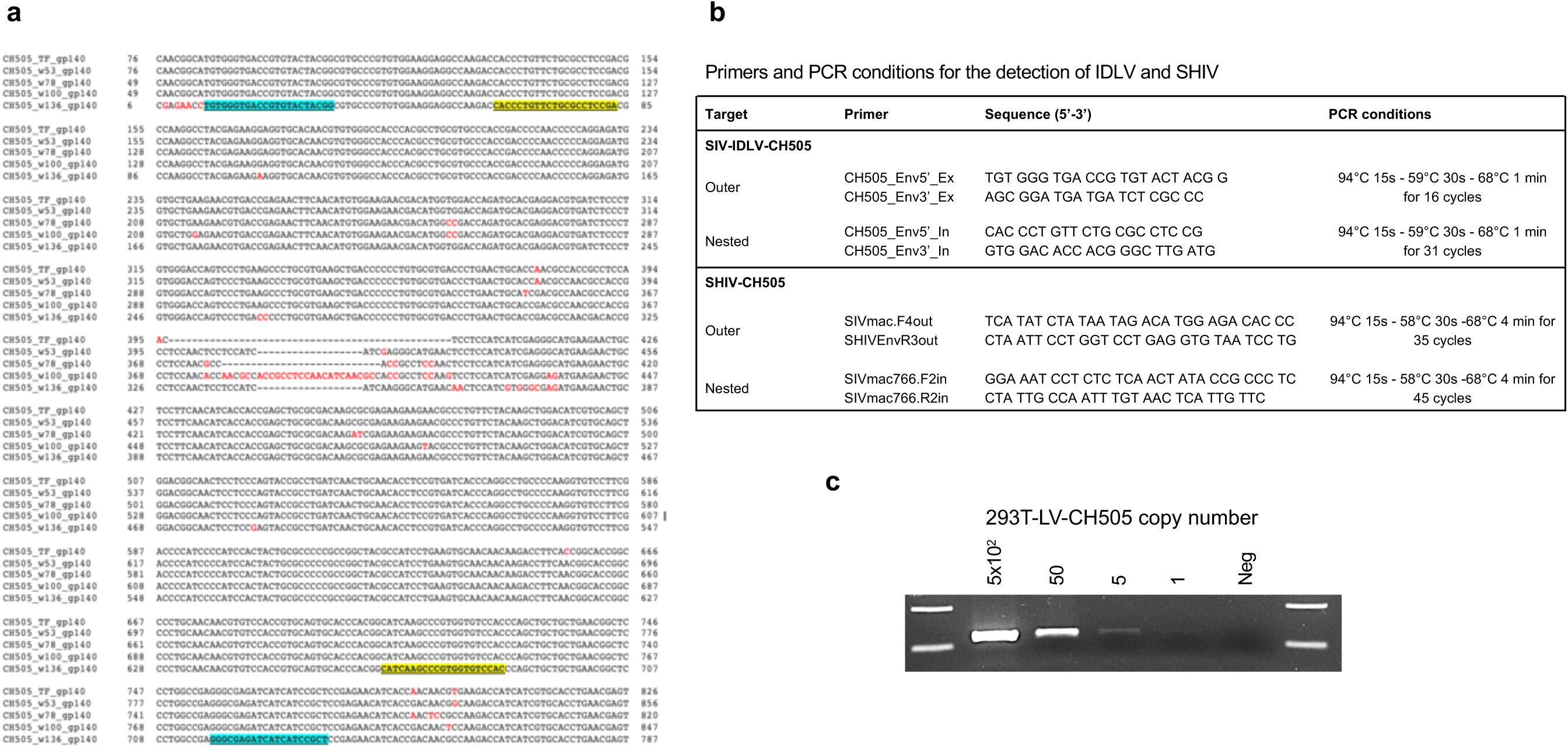
PCR primers and conditions used to detect possible mobilization and or recombination events between IDLV-CH505 and SHIV-CH505 following intravenous challenge. (a) Alignment of the CH505 sequences encoded by the IDLV-CH505 vaccines used in this study. Primer sequences are highlighted in blue for the outer primers and in yellow for the inner primers. (**b**) PCR primers and conditions used to detect IDLV-CH505 or SHIV-CH505 sequences. (**c**) Standard curves for LV- CH505 DNA were generated using serial dilutions of genomic DNA extracted from 293T cells transduced with an integrase competent lentiviral vector expressing the CH505 T/F gp140 envelope (293T-LV-CH505). Neg = genomic DNA from non-transduced cells.

## Acknowledgements

The authors thank Nancy Miller (NIAID) for her valuable input and discussion; Mark Lewis, Alex Granados and Chantelle Baker (Bioqual, Inc.) for their support with NHPs immunization and sampling; Arthur McMillan for technical assistance with binding assays; This work was supported by the National Institute of Allergy and Infectious Diseases (NIAID) #1P01AI110485-01A1; and the Non-Human Primate Central Immunology Laboratory Contract (#HHSN27201100016C) - National Institute of Allergy and Infectious Diseases (NIAID) - National Institutes of Health (NIH) - Division of AIDS (DAIDS). This work was supported in part by the European Union’s Horizon 2020 research and innovation program under grant agreement No. 681137 (EAVI2020) to A.C. and D.N.

## Author contributions

M.B. contributed to study design, oversaw the planning of the project, analyzed the data and wrote the manuscript. E.J.B. performed experiments and data analysis, including IDLV vector preparations and ELISAs. H.S. performed experiments and data analysis, including SGAs on PBMC and LNs samples. C.L. designed and analyzed the neutralization assays and provided input on study design. W.S. and Y.F. performed statistical analysis. B.B. contributed to study coordination. M.B., M.G., C. A. coordinated NHPs samples processing and distribution and performed IFN-γ ELISPOT assays. X.S. designed and analyzed epitope mapping data, prepared data visualizations, provided input on data interpretation, and edited the manuscript. B.F.H. K.S. M.A.M., D.N. A.C. contributed to study design, data analysis and editing of the manuscript. S.S. contributed to study design, oversaw ELISPOT assays, data analysis and editing of the manuscript. M.E.K. oversaw the planning and direction of the project including analysis and interpretation of the data and editing of the manuscript.

## References

1. Mascola, JR, and Haynes, BF (2013). HIV-1 neutralizing antibodies: understanding nature’s pathways. Immunol Rev 254: 225–244.

2. Williams, WB, Liao, HX, Moody, MA, Kepler, TB, Alam, SM, Gao, F, et al. (2015). HIV-1 VACCINES. Diversion of HIV-1 vaccine-induced immunity by gp41-microbiota cross-reactive antibodies. Science 349: aab1253.

3. Haynes, BF, Gilbert, PB, McElrath, MJ, Zolla-Pazner, S, Tomaras, GD, Alam, SM, et al. (2012). Immune-correlates analysis of an HIV-1 vaccine efficacy trial. N Engl J Med 366: 1275–1286.

4. Sanders, RW, van Gils, MJ, Derking, R, Sok, D, Ketas, TJ, Burger, JA, et al. (2015). HIV-1 VACCINES. HIV-1 neutralizing antibodies induced by native-like envelope trimers. Science 349: aac4223.

5. Liao, HX, Lynch, R, Zhou, T, Gao, F, Alam, SM, Boyd, SD, et al. (2013). Co-evolution of a broadly neutralizing HIV-1 antibody and founder virus. Nature 496: 469–476.

6. Doria-Rose, NA, Schramm, CA, Gorman, J, Moore, PL, Bhiman, JN, DeKosky, BJ, et al. (2014). Developmental pathway for potent V1V2-directed HIV-neutralizing antibodies. Nature 509: 55–62.

7. Gao, F, Bonsignori, M, Liao, HX, Kumar, A, Xia, SM, Lu, X, et al. (2014). Cooperation of B cell lineages in induction of HIV-1-broadly neutralizing antibodies. Cell 158: 481–491.

8. Robb, ML, Rerks-Ngarm, S, Nitayaphan, S, Pitisuttithum, P, Kaewkungwal, J, Kunasol, P, et al. (2012). Risk behaviour and time as covariates for efficacy of the HIV vaccine regimen ALVAC-HIV (vCP1521) and AIDSVAX B/E: a post-hoc analysis of the Thai phase 3 efficacy trial RV 144. Lancet Infect Dis 12: 531–537.

9. Rossi, A, Michelini, Z, Leone, P, Borghi, M, Blasi, M, Bona, R, et al. (2014). Optimization of mucosal responses after intramuscular immunization with integrase defective lentiviral vector. PLoS One 9: e107377.

10. Gallinaro, A, Borghi, M, Bona, R, Grasso, F, Calzoletti, L, Palladino, L, et al. (2018). Integrase Defective Lentiviral Vector as a Vaccine Platform for Delivering Influenza Antigens. Front Immunol 9: 171.

11. Negri, D, Blasi, M, LaBranche, C, Parks, R, Balachandran, H, Lifton, M, et al. (2016). Immunization with an SIV-based IDLV Expressing HIV-1 Env 1086 Clade C Elicits Durable Humoral and Cellular Responses in Rhesus Macaques. Mol Ther 24: 2021–2032.

12. Blasi, M, Negri, D, LaBranche, C, Alam, SM, Baker, EJ, Brunner, EC, et al. (2018). IDLV-HIV-1 Env vaccination in non-human primates induces affinity maturation of antigen-specific memory B cells. Commun Biol 1: 134.

13. Cousin, C, Oberkampf, M, Felix, T, Rosenbaum, P, Weil, R, Fabrega, S, et al. (2019). Persistence of Integrase-Deficient Lentiviral Vectors Correlates with the Induction of STING-Independent CD8(+) T Cell Responses. Cell Rep 26: 1242–1257 e1247.

14. Naldini, L, Blomer, U, Gallay, P, Ory, D, Mulligan, R, Gage, FH, et al. (1996). In vivo gene delivery and stable transduction of nondividing cells by a lentiviral vector. Science 272: 263–267.

15. Bonsignori, M, Zhou, T, Sheng, Z, Chen, L, Gao, F, Joyce, MG, et al. (2016). Maturation Pathway from Germline to Broad HIV-1 Neutralizer of a CD4-Mimic Antibody. Cell 165: 449–463.

16. Li, H, Wang, S, Kong, R, Ding, W, Lee, FH, Parker, Z, et al. (2016). Envelope residue 375 substitutions in simian-human immunodeficiency viruses enhance CD4 binding and replication in rhesus macaques. Proc Natl Acad Sci U S A 113: E3413–3422.

17. Bar, KJ, Coronado, E, Hensley-McBain, T, O’Connor, MA, Osborn, JM, Miller, C, et al. (2019). Simian-Human Immunodeficiency Virus SHIV.CH505 Infection of Rhesus Macaques Results in Persistent Viral Replication and Induces Intestinal Immunopathology. J Virol 93.

18. Coler, RN, Bertholet, S, Moutaftsi, M, Guderian, JA, Windish, HP, Baldwin, SL, et al. (2011). Development and characterization of synthetic glucopyranosyl lipid adjuvant system as a vaccine adjuvant. PLoS One 6: e16333.

19. Sanders, RW, Derking, R, Cupo, A, Julien, JP, Yasmeen, A, de Val, N, et al. (2013). A next-generation cleaved, soluble HIV-1 Env trimer, BG505 SOSIP.664 gp140, expresses multiple epitopes for broadly neutralizing but not non-neutralizing antibodies. PLoS Pathog 9: e1003618.

20. Sanders, RW, and Moore, JP (2017). Native-like Env trimers as a platform for HIV-1 vaccine design. Immunol Rev 275: 161–182.

21. LaBranche, CC, Henderson, R, Hsu, A, Behrens, S, Chen, X, Zhou, T, et al. (2019). Neutralization-guided design of HIV-1 envelope trimers with high affinity for the unmutated common ancestor of CH235 lineage CD4bs broadly neutralizing antibodies. PLoS Pathog 15: e1008026.

22. Williams, WB, Zhang, J, Jiang, C, Nicely, NI, Fera, D, Luo, K, et al. (2017). Initiation of HIV neutralizing B cell lineages with sequential envelope immunizations. Nat Commun 8: 1732.

23. Saunders, KO, Verkoczy, LK, Jiang, C, Zhang, J, Parks, R, Chen, H, et al. (2017). Vaccine Induction of Heterologous Tier 2 HIV-1 Neutralizing Antibodies in Animal Models. Cell Rep 21: 3681–3690.

24. Julien, JP, Lee, JH, Ozorowski, G, Hua, Y, Torrents de la Pena, A, de Taeye, SW, et al. (2015). Design and structure of two HIV-1 clade C SOSIP.664 trimers that increase the arsenal of native-like Env immunogens. Proc Natl Acad Sci U S A 112: 11947–11952.

25. Ringe, RP, Ozorowski, G, Rantalainen, K, Struwe, WB, Matthews, K, Torres, JL, et al. (2017). Reducing V3 Antigenicity and Immunogenicity on Soluble, Native-Like HIV-1 Env SOSIP Trimers. J Virol 91.

26. Wiehe, K, Bradley, T, Meyerhoff, RR, Hart, C, Williams, WB, Easterhoff, D, et al. (2018). Functional Relevance of Improbable Antibody Mutations for HIV Broadly Neutralizing Antibody Development. Cell Host Microbe 23: 759–765 e756.

27. Haynes, BF, Burton, DR, and Mascola, JR (2019). Multiple roles for HIV broadly neutralizing antibodies. Sci Transl Med 11.

28. Kwong, PD, and Mascola, JR (2018). HIV-1 Vaccines Based on Antibody Identification, B Cell Ontogeny, and Epitope Structure. Immunity 48: 855–871.

29. Saunders, KO, Wiehe, K, Tian, M, Acharya, P, Bradley, T, Alam, SM, et al. (2019). Targeted selection of HIV-specific antibody mutations by engineering B cell maturation. Science 366.

30. Barouch, DH, Tomaka, FL, Wegmann, F, Stieh, DJ, Alter, G, Robb, ML, et al. (2018). Evaluation of a mosaic HIV-1 vaccine in a multicentre, randomised, double-blind, placebo-controlled, phase 1/2a clinical trial (APPROACH) and in rhesus monkeys (NHP 13-19). Lancet 392: 232–243.

31. Stephenson, KE, Wegmann, F, Tomaka, F, Walsh, SR, Tan, CS, Lavreys, L, et al. (2020). Comparison of shortened mosaic HIV-1 vaccine schedules: a randomised, double-blind, placebo-controlled phase 1 trial (IPCAVD010/HPX1002) and a preclinical study in rhesus monkeys (NHP 17-22). Lancet HIV.

32. Blasi, M, and Fouda, GG (2020). Shortening HIV vaccine regimens to achieve high coverage. Lancet HIV.

33. Steichen, JM, Lin, YC, Havenar-Daughton, C, Pecetta, S, Ozorowski, G, Willis, JR, et al. (2019). A generalized HIV vaccine design strategy for priming of broadly neutralizing antibody responses. Science 366.

34. Vargas, J, Jr., Gusella, GL, Najfeld, V, Klotman, ME, and Cara, A (2004). Novel integrase-defective lentiviral episomal vectors for gene transfer. Hum Gene Ther 15: 361–372.

35. Shen, X, Duffy, R, Howington, R, Cope, A, Sadagopal, S, Park, H, et al. (2015). Vaccine-Induced Linear Epitope-Specific Antibodies to Simian Immunodeficiency Virus SIVmac239 Envelope Are Distinct from Those Induced to the Human Immunodeficiency Virus Type 1 Envelope in Nonhuman Primates. J Virol 89: 8643–8650.

36. Montefiori, DC (2009). Measuring HIV neutralization in a luciferase reporter gene assay. Methods Mol Biol 485: 395–405.

37. Tamura, K, Stecher, G, Peterson, D, Filipski, A, and Kumar, S (2013). MEGA6: Molecular Evolutionary Genetics Analysis version 6.0. Mol Biol Evol 30: 2725–2729.

